# Phylogenetic analysis of HIV-1 archived DNA in blood and gut-associated lymphoid tissue in patients receiving antiretroviral therapy: a study from Provir/Latitude45 project

**DOI:** 10.1101/2020.05.05.078634

**Authors:** Patricia Recordon-Pinson, Camille Tumiotto, Pantxika Bellecave, Franck Salin, Patricia Thebault, Annie Gosselin, Petronela Ancuta, Marcelo Soares, Jean-Pierre Routy, Hervé Fleury

## Abstract

One of the approaches to cure HIV is the use of therapeutic vaccination. We have launched the Provir/Latitude 45 study to identify conserved CTL epitopes in archived HIV-1 DNA according to the HLA class I alleles in aviremic patients under antiretroviral therapy (ART). A HIV-1 polypeptidic therapeutic vaccine based on viral sequence data obtained from circulating blood was proposed; here, our aim was to compare the proviral DNA in blood and gut-associated lymphoid tissue (GALT) at two different levels : nucleotide sequences and potential CTL epitopes. The reverse transcriptase was sequenced in both compartments using next generation sequencing (NGS) in samples from nine individuals, two of which had also single genome sequencing (SGS) performed; phylogenetic trees were established and compared; CTL epitopes were also identified according to their potential affinity for the HLA alleles. The proviral sequences of both compartments intra-patient exhibited a very low genetic divergence while it was possible to differentiate the sequences inter-patients; SGS analysis of two couples of samples confirmed that there was not a compartmentalization of the sequences intra-patient.When we simulated the CTL epitopes which can be presented by the corresponding HLA alleles in both compartments, no significant difference was observed. We conclude that the proviral DNA sequences in blood and GALT are similar and that the epitope analysis in blood can be considered as relevant to that observed in the GALT, a hard-to-reach major compartment, and can therefore be used for therapeutic vaccine approaches.

## Introduction

HIV-1 infection can be treated with ART, leading to the control of viral replication and improving the health of people living with HIV (PLWH). However, ART cannot be interrupted since this would lead to a rebound of viral replication [1,2] as virus establishes cellular (latently infected resting CD4+ memory T cells) and anatomical reservoirs very early during infection [3-8]. Gut Associated Lymphoid Tissue (GALT) is considered to be one of the main reservoirs of SIV and HIV infection [9-11]. Cure strategies for HIV-1 include therapeutic vaccination [12], although immune response observed was not able to control viral replication after ART discontinuation [13]. In this context, we launched the Provir/Latitude 45 project to identify conserved CTL epitopes in the proviral HIV-1 DNA of patients with long-term ART. The study involves *in silico* modeling based on the HIV-1 proviral DNA sequences, the HLA alleles and the HIV-1 CTL epitopes recorded in the Los Alamos database and the Immune Epitope database (IEDB) simulator following sequencing of the archived DNA from peripheral blood mononuclear cells (PBMC), i.e. from circulating blood. Since our initial work was based on proviral DNA in PBMC, we assessed whether our observations would be the same in another compartment, namely GALT. Herein, we present a phylogenetic comparison of the archived HIV-1 DNA in PBMC and GALT from patients at success of ART, together with an evaluation of theoretically conserved CTL epitopes in both compartments.

## Results

### Phylogenetic trees of intra-patient and inter-patient samples

GALT and PBMC proviral DNA samples obtained from nine patients were sequenced by NGS. The phylogenetic trees of all samples are presented in Figure 1; NGS intra-patient sequences from blood and GALT compartments analyses exhibit a low genetic divergence and are located on the same branch. When analyzing simultaneously the data from two patients (5A and 10) (Figure2) we are able, as expected, to differentiate viral sequences originating from the two patients. As a slight discrimination between PBMC and GALT samples was observed when considering NGS sequences, we carried out a SGS analysis of samples 5A and 10 (PBMC and GALT) to check a potential artefactual role of the amplification process on the discrimination between both compartments. As shown in Figure 2, this analysis indicates that there was a true intermingling of the clonal sequences for patient 5A, evidencing lack of compartmentalization. For patient 10, the clonal PBMC sequences were located at the origin of the GALT NGS part of the tree (please refer to Figure 1), then found in the PBMC part of the NGS tree while GALT clonal sequences were located at the end of the GALT tree and the origin of the PBMC tree; as for patient 5A we can conclude that there is no evidence of compartmentalization.

**Figure 1 :**
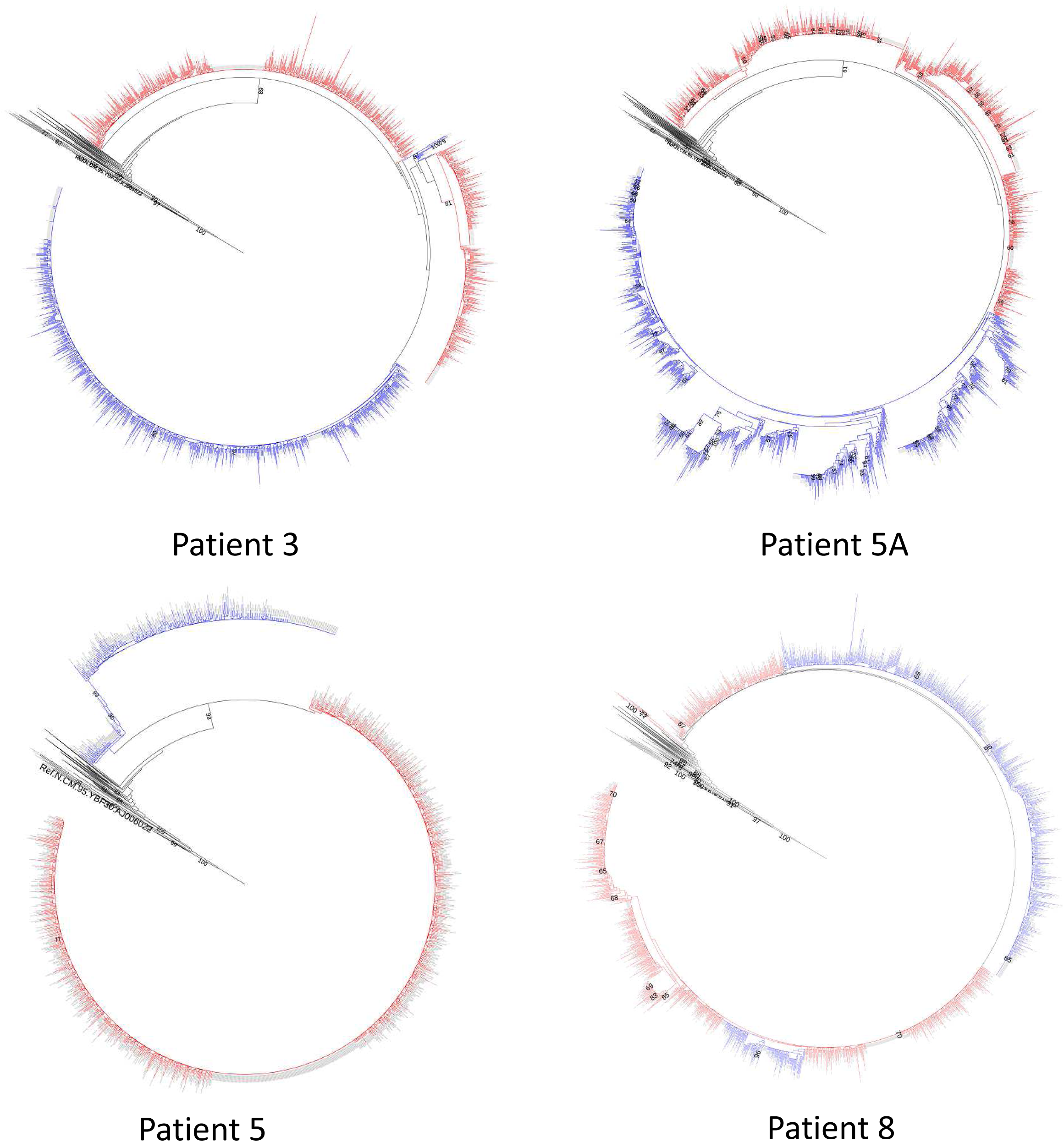
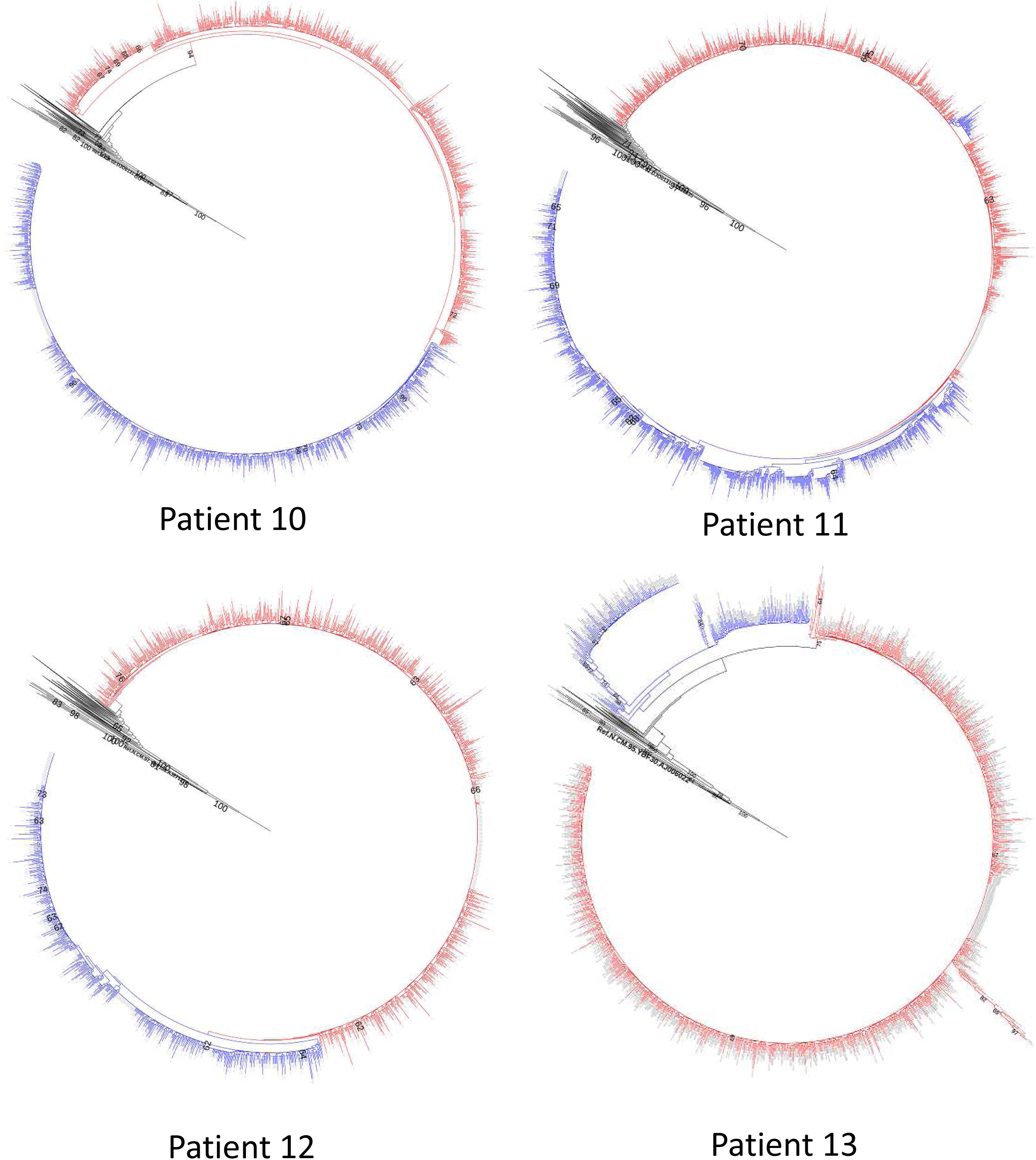
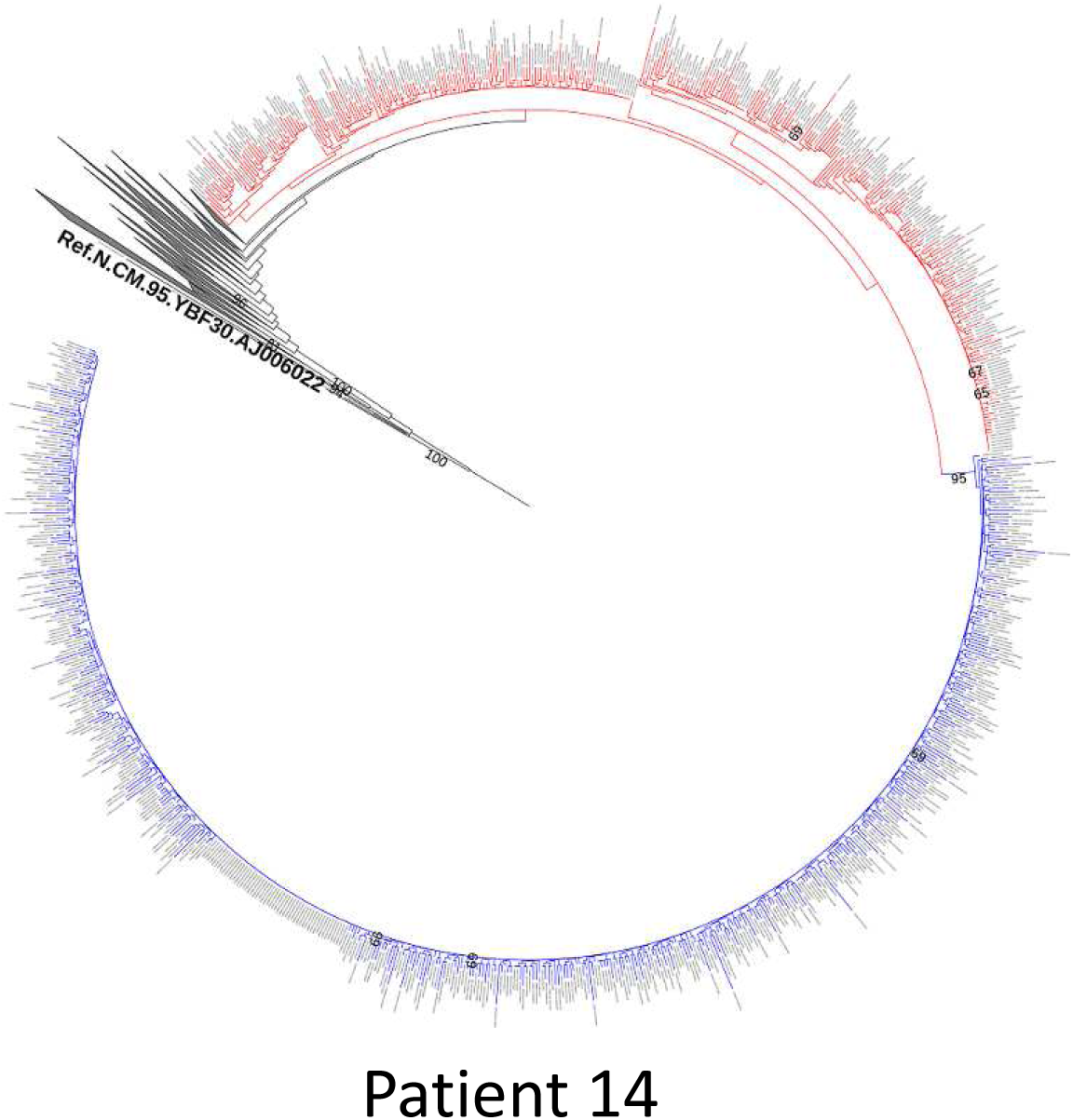
Phylogenetic trees of the NGS sequences amplified from HIV-1 proviral DNA extracted from PBMC (blue) and GALT (red) samples for all the patients. The same reference sequences were used for all trees and are notified in black. The trees are rooted on N sequences. The boostrap values greater than 50 are indicated on the branches.

**Figure 2 :**
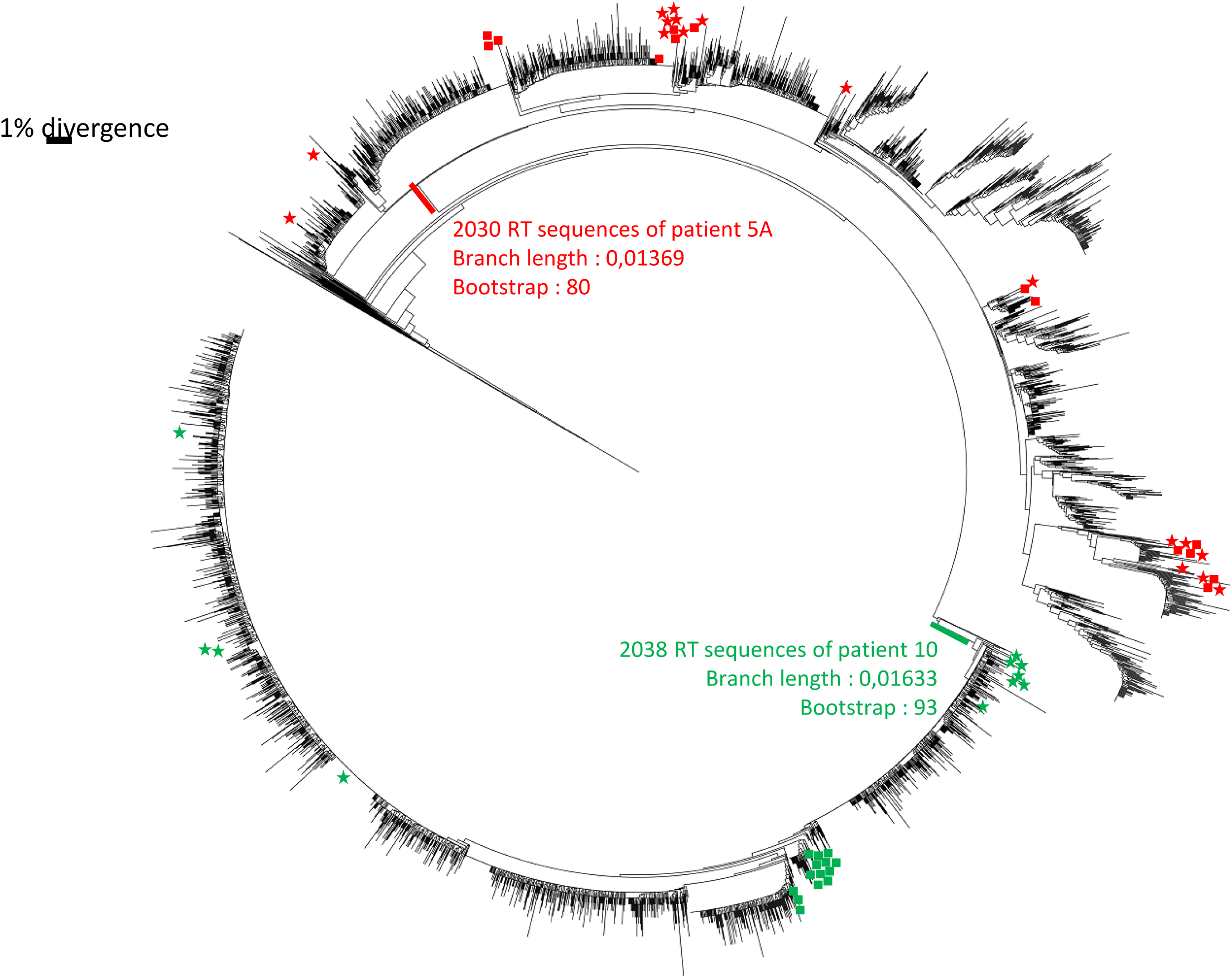
Phylogeny of RT sequences obtained by NGS and SGS from PBMC and GALT of patients 10 and 5A. The NGS sequences are in black and not detailed in PBMC or GALT (please refer to Figure1). Symbols denote sampling location and patients : patient 5A PBMC (red star), patient 5A GALT (red square), patient 10 PBMC (green star), patient 10 GALT (green square)

To confirm that all the sequences were clustered by patient, we estimated the evolutionary divergence between sequences (Table 1) considering patients 5A and 10. HIV-1 clusters were identified at maximum genetic distances between 4.5% and 7.5% and bootstrap support threshold varied between 70% and 99% [14]. As sequences from patients 5A and 10 are assembled with a median divergence of 5.3% and 2.2% respectively with bootstrap values of 80% and 93%, we confirm that these sequences from GALT and PBMC formed a specific cluster per patient.

**Table1:**
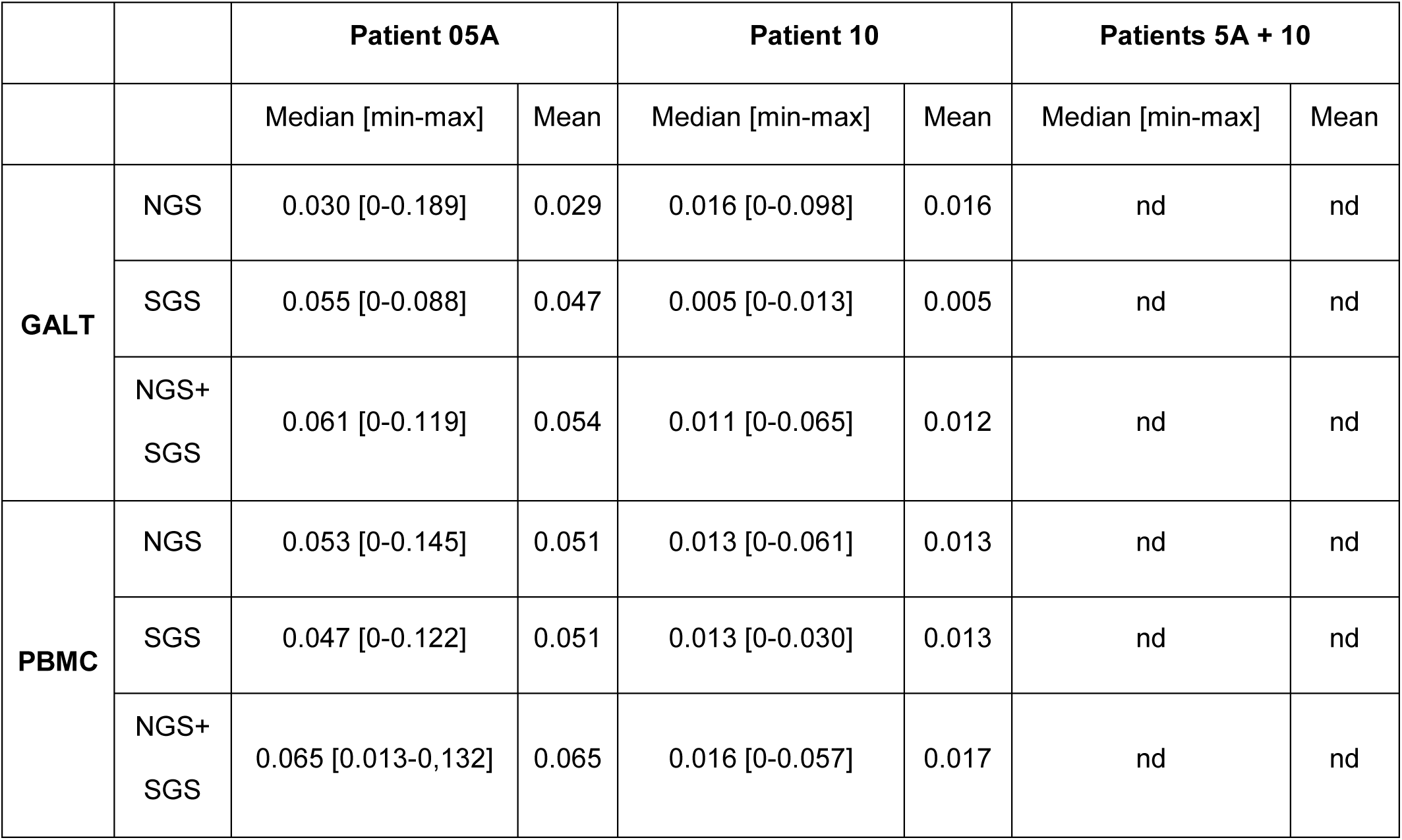
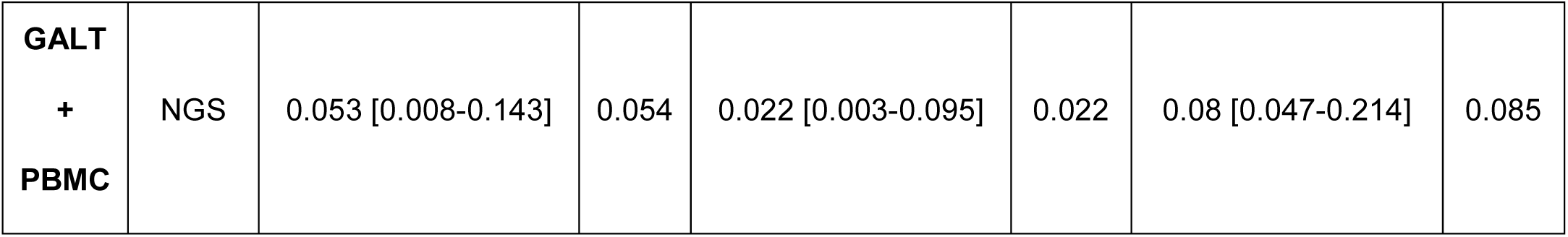
Estimates of evolutionary divergence between patients’ HIV proviral DNA sequences. Evolutionary divergences are based on the number of base substitutions per site among RT NGS and/or SGS sequences obtained from GALT and/or PBMC proviral DNA of patients 5A and 10. nd: not done

### CTL epitopes with affinity for HLA alleles

The epitopes selected by our simulation pipeline have a high theoretical affinity for the HLA alleles of the patients. We compared the inhibitory concentration 50 (IC_50_) values for each epitope based on its PBMC and GALT proviral sequences. An example is presented in Table 2. In the case of the RT (18-26) epitope, there was one variant in GALT and two in PBMC with a conserved very low IC_50_, making it possible to predict a high affinity for the B*08:01 allele. For the second epitope RT (158-166), there were two variants in both compartments. All variants were different with one in GALT at 44.89% (AIFQSSMTQ) exhibiting an IC_50_ largely>500 nanomolar (nM) (2203.8nM) and therefore having no affinity for the A*03:01 allele.

**Table 2 :** Examples of RT epitopes with theoretical affinity for the HLA alleles of the patients. Positions precise RT amino-acids involved. The numbers of reads and their percentage (%) relative to the total number of reads covering the targeted region are related to NGS results. The Inhibitory concentrations IC_50_ (nM) that reflect the affinity of the epitopes for the HLA alleles-were automatically calculated using TutuGenetics software. The modified amino-acids in the epitopes versus the reference sequence from the Los Alamos database are in bold character and underlined. When > 500 nM the IC_50_ is noted in bold italic.

| Position | HLA allele | Compartment | Epitope<br>sequence | Number<br>of reads | Reads (%) | IC<sub>50</sub> (nM) |
| --- | --- | --- | --- | --- | --- | --- |
| RT(18-26) | HLA-B*08:01 | GALT | GPKVKQWPL | 78287 | 87.73% | 41.5 |
|  |  | PBMC | GPKVKQWPL | 11562 | 87.66% | 41.5 |
|  |  |  | GPK<u>A</u>KQWPL | 233 | 1.77% | 50.5 |
| RT(158-166) | HLA-A*03:01 | GALT | AIFQSSMTK | 48154 | 44.22% | 12.8 |
|  |  |  | AIFQSSMT<u>Q</u> | 48889 | 44.89% | **2203.8** |
|  |  | PBMC | AIFQSSM<u>I</u>K | 30329 | 84.62% | 13.9 |
|  |  |  | AIF<u>R</u>SSM<u>I</u>K | 444 | 1.24% | 8.7 |

Combining all the data for the nine patients of the study and taking into account their various HLA I alleles A and B plus the obtained DNA sequences in both compartments, we found that 23 epitopes had affinity for the following alleles: A*02:01, A*03:01, A*11:01, B*08:01, B*35:01, B*40:01, B*07:02 (all data available in Table S1). The number of variants per compartment ranged from one to three. There was no significant difference in the number of variants between GALT and PBMC (Wilcoxon test p-value=0.6239). The number of epitopes exhibiting a variant with an IC_50_ >500nM was limited to one sample (please refer to the example above and to Table S1, so the variability in the compartments may not be associated with a defect on presentation of the epitopes by the alleles.

## Discussion

Archived viral DNA is found in intestinal tissue at a higher concentration than in PBMCs in ART patients [15], although the distribution of DNA in CD4+ CCR7+, transitional memory and effector memory T cells is different in blood and intestinal compartments [16]. GALT is therefore a compartment of major importance in the pathophysiology of HIV infection [17,18]. The data presented here are not related to the quantification of archived DNA but rather to a comparison of the RT sequences obtained by NGS in GALT and PBMC. In that regard, Van Marle et al. [19] have studied biopsies from infected untreated individuals and sequenced the nef and RT genes of the viral RNA from blood (PBMC) and different parts of the gut by cloning and Sanger sequencing; they concluded that there is a compartmentalization of the virus in the gut reservoir. On the other hand, Lerner et al [20] found a low diversity of the GALT and PBMC viruses in patients having experienced a voluntary treatment interruption while Imamichi et al [21] did not demonstrate any difference between RNA and DNA sequences from gut and blood of patients chronically infected with HIV-1. Studying HIV-1 infected patients at early and chronic infection stages, Rozera et al [22] found a more pronounced compartmentalization of proviral quasispecies in gut compared with PBMC samples in patients with early infection compared with chronic patients. The loss of gut/PBMC compartmentalization in more advanced stages of HIV infection was confirmed by longitudinal observation.

Regarding ART treated patients, Evering et al.[23] have studied the variability of the proviral DNA in the gut and blood compartments by SGS of the env part of the virus. They showed absence of evolution of the env sequences in the GALT and in PBMC; the authors mention that they cannot rule out the possibility of evolution in other viral genomic regions of HIV-1 such as pol which have not been investigated. Josefsson et al [24] have compared the HIV DNA in PBMC and GALT from patients being on successful ART; they have used SGS technology and showed that there is no significant difference between the sequences from both compartments. They concluded that the HIV reservoir is stable on long-term suppressive ART and raise the hypothesis that the population of infected cells exhibiting a low variability of the virus could be maintained by homeostatic cell proliferation.

The patients of our study are similar to those of the two publications mentioned above; they were long term ART-treated patients with controlled viral load and therefore a stable viral reservoir; the phylogenetic inferences obtained after NGS evidenced a very low genetic distance between the GALT and PBMC compartments intra-patient. On the other hand, it is possible to differentiate the GALT/PBMC sequences inter-patients; the SGS analysis performed for two samples, 5A and 10, plus the genetic divergence values after NGS and SGS are concordant with a high similarity between proviruses intra-patient. It must be underlined that the SGS technique decreases taq-induced recombination and nucleotide misincorporation, providing therefore a more reliable conclusion than conventional cloning [33].

Among the limitations of our study we must note the fact that only the RT part of the proviral DNA has been considered and also that we have analyzed global archived DNA molecules without differentiating noninfectious and replication competent genomes [34]; however, more recent data show that even defective proviral DNA molecules can be expressed and yield viral proteins recognized by CTL T CD8+ lymphocytes [35].

Provir/Latitude 45 project is based on the identification of conserved epitopes in the provirus with potential affinity for the dominant alleles of the population; therefore, more than 200 patients at successful ART have been investigated for these conserved epitopes [27] and a peptide cocktail has been proposed; it must be noted that this project is based on the study of the blood compartment and the question has been raised whether another compartment could yield different results; as the GALT is a major reservoir, we have then focused on a simulation of the CTL epitopes of the nine patients in both compartments with a high affinity for the corresponding HLA I alleles; having determined viral CTL epitopes in the blood compartment that should have a high affinity for the corresponding HLA A and B alleles of the patients individually, we have raised the question of a full identity or not of these epitopes in the gut compartment; a significant difference would be a hurdle for the vaccine strategy based on the blood analysis; clearly, however, the differences between the two compartments are slight even if there are variants at the epitope locations (but with a close affinity for the HLA alleles); the global choice of the epitopes at the blood level seems therefore extrapolable to the GALT level. In conclusion, our results confirm that the proviruses in GALT and PBMC are very similar in these patients having experienced a long-term ART success and who could be the target population of choice for a therapeutic vaccine. They also provide new data on the high similarity of the CTL epitopes in both compartments according to their potential affinity for the HLA alleles of the patients investigated. Taken together, these results indicate that the analysis of the blood compartment can provide results that can extrapolated to the gut compartment, a major reservoir of HIV. The strategies used in the future, whatever they might be, will have to eradicate the virus from the blood and also from the different tissue reservoirs including the gut.

## Materials and Methods

### Study participants

A total of nine PLWH were recruited at MUHC. Cells were isolated at the CHUM Research Centre, Montreal, Quebec, Canada. All participants were MSM infected with HIV-1 subtype B. They were under successful ART which had been initiated between at least one year up to 14 years after HIV infection, while biopsies were obtained between four to 26 years after ART initiation; the main HLA alleles of the series investigated were representative of a Caucasian population (among them HLA A*02:01 and B*07:02); detailed clinical information on study cohort participants has been previously published [28].

### Sigmoid colon biopsies and blood cells

Sigmoid biopsies (≈32 biopsies/donor) were collected from HIV-infected individuals receiving ART during colonoscopy and processed using Liberase DL (Roche Diagnostics), as previously described (28). Matched peripheral blood (20 ml/donor) was collected on the same day from biopsy donors and immediately processed with Ficoll for PBMC isolation, performed in parallel with cell extraction from biopsy tissue.

### Extraction of total HIV DNA

Cells from gut biopsies and PBMC were suspended in 350 μL of RLT buffer with β-mercaptoethanol and total DNA was extracted using the Qiagen AllPrep DNA/RNA kit. The extracted DNA was then sent to Bordeaux University Hospital for further investigation.

### Identification of HLA alleles

The method used for the molecular characterization of HLA alleles A and B in the Montreal cohort has been described previously [25]. Briefly, intermediate-to-high resolution sequencing of extracted DNA was performed by reverse Polymerase Chain Reaction-Sequence Specific Oligonucleotide (PCR-SSO) hybridization by using the LuminexH flow beads LabTypeH assay (InGen) for the A and B loci. Allelic ambiguities were resolved by PCR-Sequence Specific Primer (SSP) amplification by using Olerup assays. Allele assignment was performed by comparison with official nomenclature and validated by the WHO committee for HLA system factors. All HLA A and B alleles have been determined for all patients. They are presented and discussed in Table 2 and Table S1 only when there affinity for the reference epitopes exhibits an inhibitory concentration 50, IC_50_<50nM (please refer to the sub chapter Prediction of affinity between epitopes and HLA alleles using NGS data).

### NGS of HIV proviral DNA

We used the method already published [29,30] to amplify fragment B, i.e. polymerase (Pol) region including RT and integrase. After DNA extraction, nested PCR was carried out with the following outer primers: 1st F ATGATAGGGGGAATTGGAGGTTT (HXB2 loci 2388-2410), 1st R CCTGTATGCAGACCCCAATATG (5264-5243), and inner primers 2nd F GACCTACACCTGTCAACATAATTGG (2485-2509), 2nd R CCTAGTGGGATGTGTACTTCTGAACTTA (5219-5192). The PCR products were purified and quantitated. The library was prepared using the Nextera XT DNA Sample Preparation kit. A purification step was performed and the library was quantified by Tapestation technology. Each individual library was then sequenced on a MiSeq Illumina platform. Raw data (FASTQ files) were submitted to the SmartGene® NGS HIV-1 module to generate a BAM file for each patient and each sample was processed for further analysis [31]. The study was carried out using only the Pol RT part region of the sequences obtained.

### Phylogenetic analysis following NGS

Using Galaxy and Clustal software, RT gene sequences from the two compartments (GALT and PBMC) per patient were selected for neighbor-joining analysis from matrix distances calculated after gapstripping of alignments, with a Kimura two-parameter algorithm and bootstrap analysis. To do so, an alignment was generated that included only reads with lengths > 400 bp corresponding to a given region of RT (variable according to the patient, from amino-acids 39 to 202). This length limitation explains the small number of reads used for this analysis compared to the total number of reads covering this region. Phylogenetic trees were visualized using ITOL (Interactive Tree of Life) software [32].

### SGS and analysis

SGS was carried out using samples from two patients (5A and 10) according to the method of Palmer et al [33]. The total extracted DNA of both compartments was diluted in TE buffer at a dilution yielding a PCR product in 3 out of 10 PCRs. In this case, according to Poisson’s distribution, the dilution contains 1 copy of cDNA per positive PCR at about 80% of the time. Two rounds of PCR for RT amplification were followed by visualization of the PCR products by capillary electrophoresis using QIAxcel DNA screening kit. The 1:9 dilution was found to be optimal for Sanger sequencing and the sequences (assuming that there was no mixture of population) of PBMC and GALT obtained were aligned by Clustal to obtain a neighbor-joining tree.

### Prediction of affinity between epitopes and HLA alleles using NGS data

The TutuGenetics software [36] allows the automatic identification of CTL epitopes that can theoretically be presented by the HLA I alleles of the patients. It links the sequence data, the identity of the HLA alleles, the Los Alamos HIV database for CTL epitopes and the International Immune Epitope Database (IEDB) simulator. It does not only identify CTL epitopes in the LANL database but also proposes predictive epitopes as previously published [27]. Using the TutuGenetics software, the theoretical affinity between the CTL epitopes encoded in the RT region and the HLA alleles of each patient at both compartments was calculated. We focused only on those epitopes that had a theoretical affinity for a defined allele of the patient with an MHC IC_50_ <50nM. When variants were identified, they were evaluated for their IC_50_<50nM or in ranges between 50-500 nM. It should be noted that the number of reads analyzed was higher since only variants present at a frequency > 1% at a given position of the CTL epitopic peptide sequence were considered.

### Evolutionary divergence and statistical analysis

The median, mean and range of the number of base substitutions per site between RT sequences were calculated. Analyses were conducted using the Maximum Composite Likelihood model [37].The analysis involved 4100 nucleotide sequences. Codon positions included were 1st+2nd+3rd+Noncoding. All positions containing gaps and missing data were eliminated. There were a total of 376 positions in the final dataset. Evolutionary analyses were conducted in MEGA7 [38].

The Wilcoxon test was used for analysis of the number of variants at the different positions of epitopes with a theorical affinity for defined alleles within both compartments; the IC_50_ values considered were <500nM and >500nM.

## List of abbreviations

GALT: gut associated lymphoid tissue;
NGS: next generation sequencing;
PLWH: people living with HIV;
PBMC: peripheral blood mononuclear cells;
IC_50_: inhibitory concentration 50;
SGS: single genome sequencing;
Pol: polymerase;
RT: reverse transcriptase;
ART: antiretroviral therapy

## Declarations

### Ethics

this study using colon biopsies and blood from patients living with HIV (PLWH) was approved by the Research Ethics Boards of the Research Institute of McGill University Health Centre (MUHC), Montreal, QC, Canada and the Centre Hospitalier de l’Université de Montréal (CHUM) Research Centre, Montréal, QC, Canada. The study also received approval from the Comité d’Ethique du Sud Ouest et Outremer (DC 2012/48) and was conducted in accordance with the Declaration of Helsinki. Each participant provided written informed consent before any study procedure.

### Availability of data

the NGS sequences are available in GenBank under accession number PRJN557560; the Sanger sequences of the SGS study are available in GenBank under accession numbers MN250226 to MN250287

### Authors’ contributions

HF designed the study, found the main source of funding (MSD Avenir) and wrote the manuscript with the help of PP and contributions from MS, PA and JPR; JPR recruited the patients; PP, CT, PB, FS, PT, and AG executed technical parts of the study and were involved in analysis of the data. All authors read and approved the final manuscript.

## Acknowledgments

we thank Lucie Richard, an engineering student at the university of Bordeaux, for her excellent technical collaboration. The valuable advices and comments by Z Brumme (British Columbia Centre for Excellence in HIV/AIDS, Vancouver, BC, Canada) were greatly appreciated.

We would like to thank Maria Fraraccio for her administrative support in Montreal and Fredéric Perry for his administrative assistance at the DRCI CHU in Bordeaux. Ray Cooke is also thanked for copyediting.

