## Supplemental Table 1 for "Phylogenetic analysis of HIV-1 archived DNA in blood and gut-associated lymphoid tissue in patients receiving antiretroviral therapy: a study from Provir/Latitude45 project"

**Table S1. Comparison of the CTL epitopes with an affinity to the HLA alleles of the patients in both compartments, blood (PBMC) and GALT**

All CTL epitopes with a theorical affinity ($IC_{50}$ <50nM) for the HLA alleles A and B of the patients are listed in the Los Alamos database with the exception of RT 110-118, RT 110-118 and RT 240-249 which are listed in our previous work regarding HFVAC [27].

The table includes each patient's identification number, the position of the peptide in the HIV genome, the compartments (GALT or PBMC), the HLA allele with a theorical affinity for the peptide, the number of reads following NGS, the percentage of the epitope within the variants, the calculated $IC_{50}$ in nM

The amino acids which differ within the reference epitopes are noted in bold and underlined.

The value of $IC_{50}$ > 500nM is in bold and italic

| Patient | Position | compartment | HLA | Epitope | Nb of reads | % of reads | IC50 |
|---|---|---|---|---|---|---|---|
| Patient 3 | RT(73-82) | GALT | HLA-A*03:01 | KLVDFRELNK | 67342 | 85.9% | 20.3 |
| | | PBMC | | KLVDFRELNK | 11061 | 85.01% | 20.3 |
| | | | | KLVD**L**RELNK | 160 | 1.23% | 38.4 |
| | RT(107-115) | GALT | HLA-B*35:01 | TVLDVGDAY | 78213 | 89.02% | 21.8 |
| | | PBMC | | TVLDVGDAY | 11922 | 85.58% | 21.8 |
| | RT(156-164) | GALT | | SPAIFQCSM | 79033 | 65.91% | 42.8 |
| | | | | SPAIF**R**CSM | 26626 | 22.20% | 81 |
| | | PBMC | | SPAIFQCSM | 12744 | 66.62% | 42.8 |
| | | | | SPAISQCSM | 3551 | 18.56% | 62.8 |
| | RT(158-166) | GALT | HLA-A*03:01 | AIFQCSMTK | 79660 | 65.26% | 11.5 |
| | | | | AIF**R**CSMTK | 27113 | 22.21% | 7.8 |
| | | PBMC | | AIFQCSMTK | 13068 | 66.54% | 11.5 |
| | | | | AI**S**QCSMTK | 3642 | 18.54% | 36.3 |
| Patient 5 | RT(18-26) | GALT | HLA-B*08:01 | GPKVKQWPL | 22438 | 91.06% | 41.5 |
| | | PBMC | | GPKVKQWPL | 395 | 19.16% | 41.5 |
| | | | | **S**PKVKQWPL | 1475 | 71.53% | 13.7 |
| Patient 8 | RT(181-189) | GALT | HLA-A*02:01 | YQYMDDLYV | 61539 | 70.02% | 11.5 |
| | | | | YQY**I**DDLYV | 14356 | 16.33% | 19.8 |
| | | PBMC | | YQYMDDLYV | 44805 | 82.735% | 11.5 |

| | | | | | | | |
|---|---|---|---|---|---|---|---|
| | | | | YQ**C**MDDLYV | 1823 | 3.37% | 99.6 |
| | | | | YQYMD**G**LYV | 696 | 1.29% | 11.2 |
| | RT(232-241) | GALT | | YELHPDKWTV | 78253 | 85.96% | 43 |
| | | PBMC | | YELHPDKWTV | 47691 | 80.85% | 43 |
| Patient 10 | RT(73-82) | GALT | HLA-A*03:01 | KLVDFRELNK | 80167 | 88.30% | 20.3 |
| | | | HLA-A*11:01 | KLVDFRELNK | 80167 | 88.30% | 40.8 |
| | | PBMC | HLA-A*03:01 | KLVDFRELNK | 44236 | 86.85% | 20.3 |
| | | | HLA-A*11:01 | KLVDFRELNK | 44236 | 86.85% | 40.8 |
| | RT(107-115) | GALT | HLA-B*35:01 | TVLDVGDAY | 87236 | 87.42% | 21.8 |
| | | PBMC | HLA-B*35:01 | TVLDVGDAY | 41471 | 81.86% | 21.8 |
| | RT(156-164) | GALT | HLA-B*35:01 | SPAIFQSSM | 106719 | 87.67% | 43.5 |
| | | PBMC | HLA-B*35:01 | SPAIFQSSM | 59142 | 87.72% | 43.5 |
| | RT(158-166) | GALT | HLA-A*03:01 | AIFQSSMTK | 106970 | 87.17% | 12.8 |
| | | | HLA-A*11:01 | AIFQSSMTK | 106970 | 87.17% | 8.3 |
| | | PBMC | HLA-A*03:01 | AIFQSSMTK | 60074 | 87.49% | 12.8 |
| | | | HLA-A*11:01 | AIFQSSMTK | 60074 | 87.49% | 8.3 |
| | RT(240-249) | GALT | HLA-A*11:01 | TVQPIVLPEK | 4368 | 11.12% | 17.1 |
| | | | | TVQPI**M**LPEK | 29503 | 75.08% | 13 |
| | | PBMC | | TVQPI**M**LPEK | 64133 | 86.41% | 13 |
| | | | | TVQPI**M**L**L**EK | 785 | 1.06% | 13.3 |
| Patient 11 | | GALT | HLA-A*02:01 | YQYMDDLYV | 156826 | 88.35% | 11.5 |
| | | | | YQYMD**V**LYV | 3028 | 1.71% | 4.8 |

| | | | | | | | |
|---|---|---|---|---|---|---|---|
| | RT(181-189) | PBMC | | YQYMDDLYV | 59492 | 45.16% | 11.5 |
| | | | | YQY**V**DDLYV | 60939 | 46.26% | 15.9 |
| | RT(232-241) | GALT | | YELHPDKWTV | 63110 | 85.30% | 43 |
| | | | | Y**G**LHPDKWTV | 940 | 1.27% | 43 |
| | | PBMC | | YELHPDKWTV | 53564 | 88.84% | 43 |
| Patient 12 | RT(18-26) | GALT | HLA-B*08:01 | GPKVKQWPL | 78287 | 87.73% | 41.5 |
| | | PBMC | | GPKVKQWPL | 11562 | 87.66% | 41.5 |
| | | | | GPK**A**KQWPL | 233 | 1.77% | 50.5 |
| | RT(181-189) | GALT | HLA-A*02:01 | YQYMDDLYV | 131768 | 89.08% | 11.5 |
| | | PBMC | | YQY**V**DDLYV | 39724 | 87.60% | 15.9 |
| | | | | YQY**V**DDL**H**V | 544 | 1.20% | 108.4 |
| | RT(232-241) | GALT | | YELHPDKWTV | 60091 | 85.30% | 43 |
| | | PBMC | | YELHPDKWTV | 36768 | 85.98% | 43 |
| Patient 14 | RT(18-26) | GALT | HLA-B*08:01 | GPKVKQWPL | 15886 | 90.38% | 41.5 |
| | | PBMC | | GPKVKQWPL | 9533 | 87.54% | 41.5 |
| | RT(73-82) | GALT | HLA-A*03:01 | KLVDFRELNK | 31941 | 42.82% | 20.3 |
| | | | | K**I**VDFRELNK | 33393 | 44.76% | 36.5 |
| | | | | KLVD**I**RELNK | 906 | 1.22% | 35.2 |
| | | PBMC | | KLVDFRELNK | 18877 | 86.69% | 20.3 |
| | RT(156-164) | GALT | HLA-B*07:02 | SPAIFQSSM | 97283 | 89.37% | 9 |
| | | PBMC | | SPAIFQSSM | 30305 | 85.74% | 9 |
| | | | | SPAIF**R**SSM | 449 | 1.27% | 3.3 |
| | RT(158-166) | GALT | HLA-A*03:01 | AIFQSSMTK | 48154 | 44.22% | 12.8 |
| | | | | AIFQSSMT**Q** | 48889 | 44.89% | **2203.8** |
| | | PBMC | | AIFQSSM**I**K | 30329 | 84.62% | 13.9 |
| | | | | AIF**R**SSM**I**K | 444 | 1.24% | 8.7 |
| | RT(181-189) | GALT | HLA-A*02:01 | YQYMDDLYV | 40432 | 44.16% | 11.5 |
| | | | | YQY**V**DDLYV | 41705 | 45.55% | 15.9 |
| | | PBMC | | YQYMDDLYV | 31645 | 84.77% | 11.5 |
| | | | | YQYMDDL**C**V | 770 | 2.06% | 77.5 |

| | | | | | | | |
|---|---|---|---|---|---|---|---|
| | | | | YQYMDD**S**YV | 433 | 1.16% | 22.5 |
| | RT(232-241) | GALT | HLA-A*02:01 | YELHPDKWTV | 20928 | 84.55% | 43 |
| | | | | YEL**R**PDKWTV | 736 | 2.97% | 323.3 |
| | | PBMC | | YELHPDKWTV | 37449 | 83.53% | 43 |
| | | | | Y**K**LHPDKWTV | 1440 | 3.21% | 43 |
